# Molecular phylogeny of Puerto Rico Bank dwarf geckos (Squamata: Sphaerodactylidae: *Sphaerodactylus*)

**DOI:** 10.1101/2021.03.23.436310

**Authors:** R. Graham Reynolds, Aryeh H. Miller, Alejandro Ríos-Franceschi, Clair A. Huffine, Jason Fredette, Nicole F. Angeli, Sondra I. Vega-Castillo, Liam J. Revell, Alberto R. Puente-Rolón

**Author notes:** Corresponding author.; URL: http://www.caribbeanboas.org.

## Abstract

The genus *Sphaerodactylus* is a very species-rich assemblage of sphaerodactylid lizards that has undergone a level of speciation in parallel to that of the well-known *Anolis* lizards. Nevertheless, molecular phylogenetic research on this group consists of a handful of smaller studies of regional focus (e.g., western Puerto Rico, the Lesser Antilles) or large-scale analyses based on relatively limited sequence data. Few medium-scale multi-locus studies exist— for example, studies that encompass an entire radiation on an island group. Building upon previous work done in Puerto Rican *Sphaerodactylus*, we performed multi-locus sampling of *Sphaerodactylus* geckos from across the Puerto Rico Bank. We then used these data for phylogeny estimation with near-complete taxon sampling. We focused on sampling the widespread nominal species *S. macrolepis* and in so doing, we uncovered a highly divergent and morphologically distinct lineage of *Sphaerodactylus macrolepis* from Puerto Rico, Culebra, and Vieques islands, which we recognize as *S. grandisquamis* (Stejneger, 1904) on the basis of molecular and morphological characters. *S. grandisquamis* co-occurs with *S. macrolepis* only on Culebra Island but is highly genetically differentiated and morphologically distinct. *Sphaerodactylus macrolepis* is now restricted to the eastern Puerto Rico Bank, from Culebra east through the Virgin Islands and including the topotypic population on St. Croix. We include additional discussion of the evolutionary history and historical biogeography of the *Sphaerodactylus* of the Puerto Rican Bank in the context of these new discoveries.

---

Harris and Kluge (1984) describe the genus *Sphaerodactylus* as “one of the most speciose genera of gekkonid lizard”— with 106 currently recognized taxa spread throughout the Caribbean basin and adjacent continental Americas (Uetz et al., 2018). The bulk of this diversity (approximately 80 species) occurs in the West Indies (Schwartz and Henderson, 1991). *Sphaerodactylus* have diversified across the West Indies colonizing nearly every island bank in the Caribbean archipelago from large to small (Maury et al., 1990; Schwartz and Henderson, 1991). The group is especially fascinating as they appear to have undergone repeated ecological and evolutionary radiations on larger Greater Antillean islands such as the Puerto Rico Bank (Grant, 1931a; Rivero, 1998, 2006). Populations of some species can reach astonishingly high densities, and *Sphaerodactylus* includes the species with the highest recorded density of any terrestrial vertebrate taxon (*S. macrolepis* on Guana Island in the British Virgin Islands; Rodda et al., 2001).

Despite its impressive species richness and the high population density of some members of the group, Greater Antillean *Sphaerodactylus* have been surprisingly understudied over the past 25 years or so, particularly when compared to other squamate taxa of the region (e.g., *Anolis*; Losos, 2009). Some recent natural history research (e.g., Allen and Powell, 2014; de Queiroz and Losos, 2017; Kelehear et al., 2017) has begun to reveal the ecology of a few species. More is likely known regarding phylogenetic relationships than ecology, and previous work using molecular data (e.g., Haas, 1991, 1996; Thorpe et al., 2008; Díaz-Lameiro et al., 2013; Surget-Groba and Thorpe, 2013) has provided some valuable hypotheses regarding sphaerodactylid evolution in the region. This work, however, was based on a single genetic marker (mtDNA) or allozymes. A large-scale phylogenetic study of all squamates by Pyron et al. (2013) provides a phylogenetic hypothesis for examining sphaerodactylid evolutionary relationships; however, the vast majority of the gecko relationships in this tree were based exclusively on approximately five hundred base pairs (bp) of mitochondrial 16S sequence data and many species branches have low support.

We investigated the evolutionary relationships among the species of *Sphaerodactylus* on the Puerto Rico Bank (PRB) and the St. Croix Bank (SXB). This archipelago consists of the island of Puerto Rico along with the Spanish, British, and United States Virgin Islands (excluding St. Croix). Seven species in this genus are found on the PRB/SXB, while a further three species (related to PRB/SXB lineages) are found on the islands of Mona, Monito, and Desecheo (Thomas and Schwartz, 1966; Schwartz and Henderson, 1991; Rivero, 1998, 2006; Díaz-Lameiro et al., 2013) which are located west of the main island of Puerto Rico but are not part of the PRB (Fig. 1). One Puerto Rican species (*S. macrolepis*) also occurs on the SXB (the topotypic location for the species), along with at least one other taxon: the endemic *S. beattyi* (Fig. 1).

**Figure 1:**
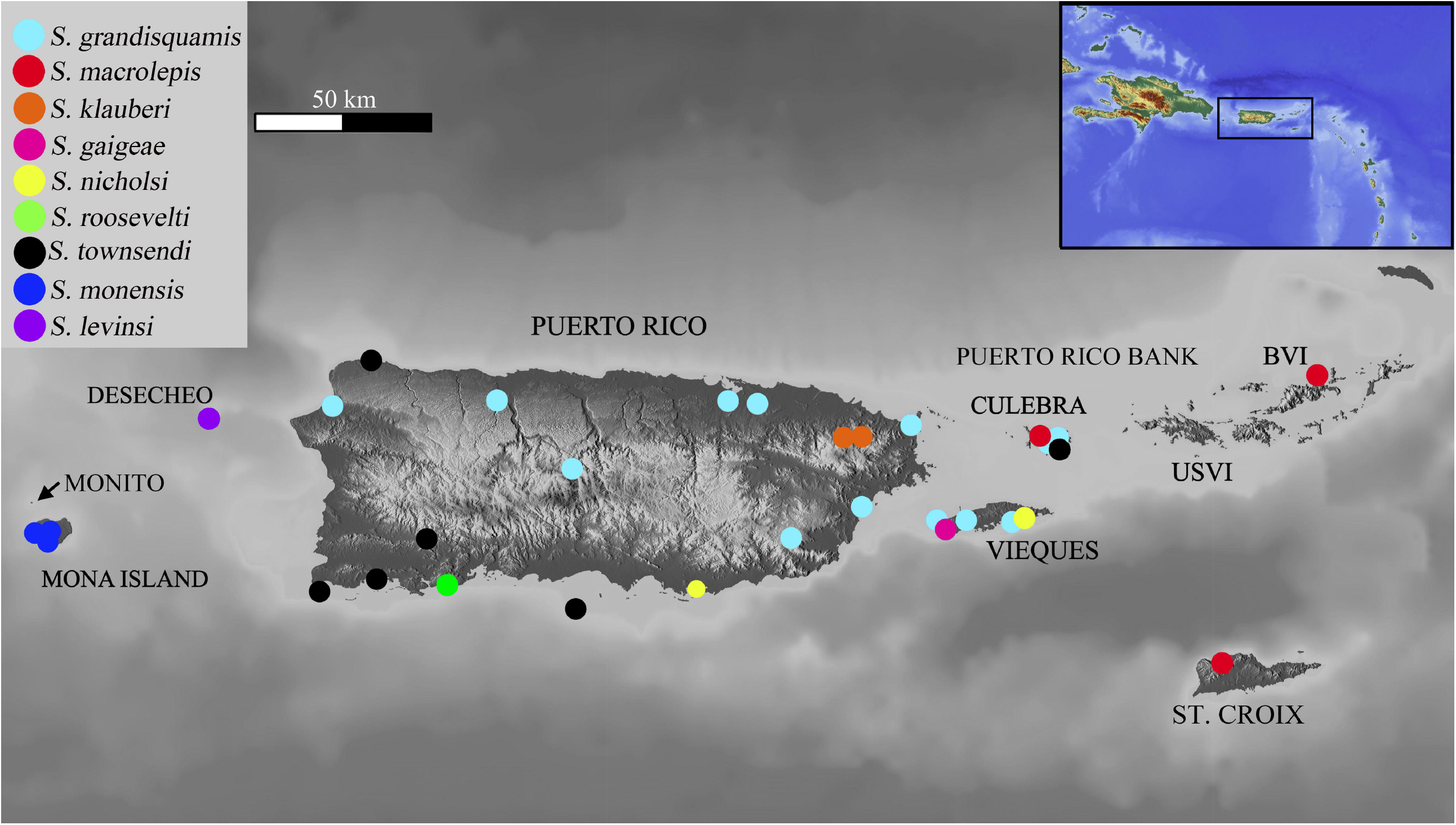
Sampling map of all *Sphaerodactylus* localities on the Puerto Rican Bank included in this study, with the northeastern Caribbean region shown in the inset. Note that this is not a complete range map for these species. Islands on Puerto Rico Bank are shown, as is St. Croix. Island elevation is shown in terrain colors, and the approximate extent of the inundated Puerto Rico Bank is shown in light blue. Islands mentioned in the text are also labeled. See Supplemental Table S1 for specimen and locality information.

Puerto Rican *Sphaerodactylyus* have been more well-studied than species from any other island in the Greater Antilles. This is particularly true of the species from western Puerto Rico and the islands of the Mona passage (e.g., Díaz-Lameiro, 2011; Díaz-Lameiro et al., 2013). In addition, Murphy et al. (1984) provided evidence of a stable hybrid zone between two nominally distinct taxa (*S. nicholsi* and *S. townsendi*) in southern Puerto Rico (Pinto et al., 2019).

Investigations of different herpetofaunal lineages have undergone biogeographic diversification in different ways on the PRB (Brandley and de Queiroz, 2004; Barker et al. 2012, 2015; Rodríguez-Robles et al., 2015; Reynolds et al. 2015, 2017). This led us to suspect that a deeper look at the phylogenetics of *Sphaerodactylus* of the PRB/SXB might also reveal new and potentially unexpected findings. A study investigating the relationships among *Sphaerodactylus* inhabiting the islands in the Mona Passage found that a cryptic lineage related to *S. klauberi* likely lives in the Rincón region of northwestern Puerto Rico (Díaz-Lameiro et al., 2013). Based upon these studies, we undertook a more detailed investigation of relationships within and among the *Sphaerodactylus* of the PRB/SXB to see if diversification has taken place within the species inhabiting this area.

We particularly focused on *S. macrolepis*— which was originally described from the SXB and is thought to also abound across the PRB (Schwartz and Henderson, 1991; Rivero, 1998). We obtained data from across the range of this species on the Puerto Rico Bank, inclusive of six of nine recognized subspecies (Rivero, 1998, 2006). We deployed Bayesian Inference (BI) and Maximum Likelihood (ML) methods to assess species-level relationships within the PRB/SXB *Sphaerodactylus* phylogeny that we generating using this data set. Our multi-locus sequence data and morphological analyses revealed a cryptic lineage of the widespread *S. macrolepis* from Culebra, Vieques, and the main island Puerto Rico, representing a heretofore unknown species of *Sphaerodactylus* which we re-describe as *S. grandisquamis* (Stejneger, 1904).

## MATERIALS AND METHODS

### Sample Collection

We sampled *Sphaerodactylus* from Culebra Island, a 27 km^2^ island located on the Puerto Rico Bank and situated approximately 30 km to the east of the main island of Puerto Rico (Fig. 1). Sampling details for Culebra Island are described in Ríos-Franceschi et al. (2016), and our collections focused on the area around Monte Resaca, located on the northcentral end of the island. We examined eight voucher specimens and 18 tissue samples that were collected as part of an initial study of the herpetofauna of Monte Resaca (Ríos-Franceschi et al., 2016), from which we obtained the DNA samples used in this study. Our collections also included 48 *Sphaerodactylus* tissue samples from across the PRB representing six of the seven species (*S. gaigeae, S. klauberi, S. macrolepis, S. nicholsi, S. roosevelti*, and *S. townsendi*), as well as species from Isla Mona (*S. monensis*) and Isla Desecheo (*S. levinsi*) west of the Puerto Rico Bank (Fig. 1). The only PRB species not included in our tissue dataset is *S. parthenopion*, which is endemic to the British Virgin Islands east of Culebra. Additional samples for populations within *S. macrolepis* included *S. macrolepis inigoi* (n = 12) from Vieques, *S. m. guarionex* (n = 4; Fig. 2, F), *S. m. grandisquamis* (n = 5), and *S. m. spanius* (n = 1) from the main island of Puerto Rico, and *S. m. macrolepis* (n=4) from Guana Island, British Virgin Islands. To obtain these tissues, we captured wild specimens via diurnal and nocturnal hand-capture and removal of 5 mm of the tail, which we stored in 95% EToH. We supplemented these with museum loans of liver tissue subsamples (Table S1).

**Figure 2.**
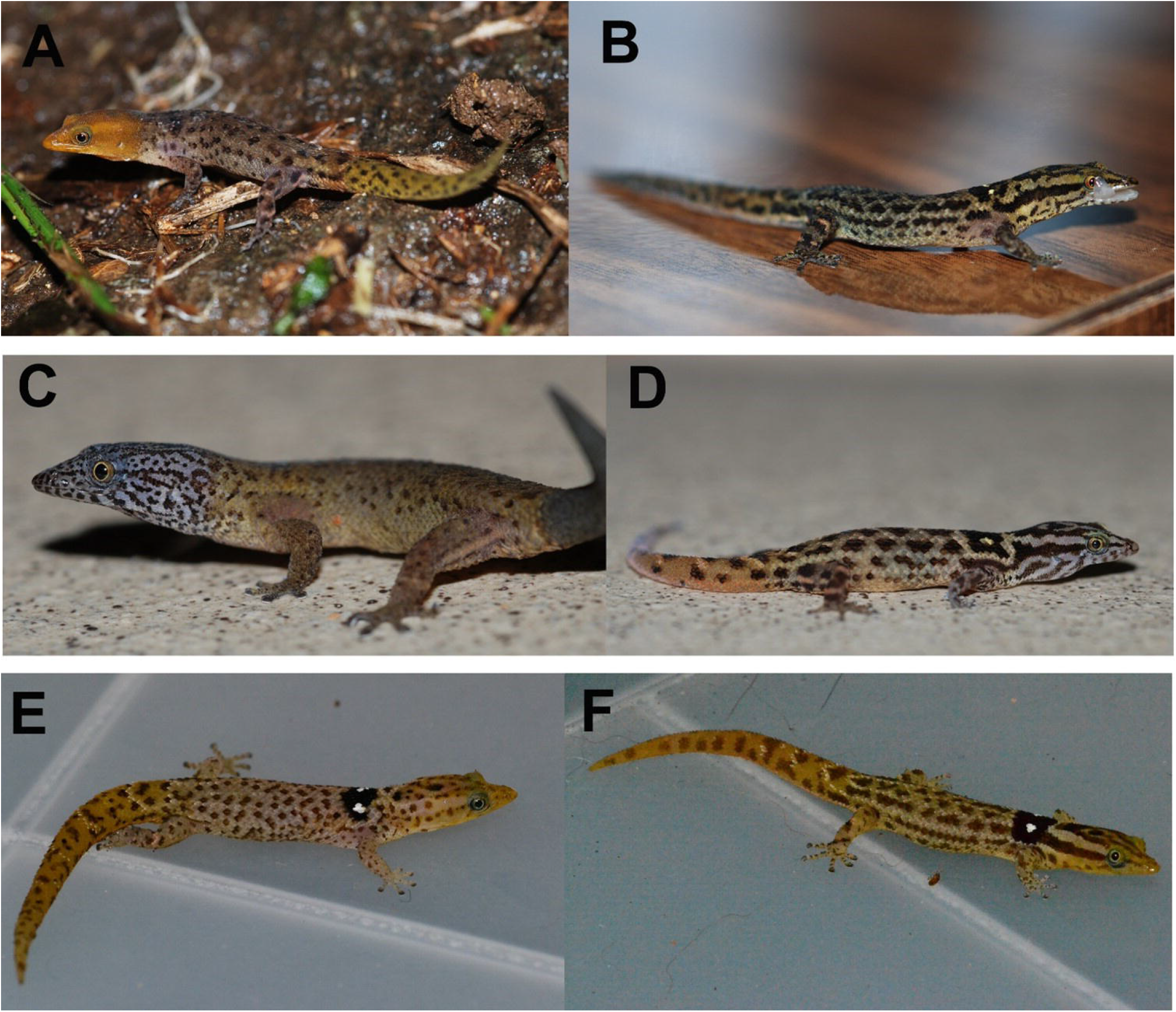
*Sphaerodactylus grandisquamis inigoi* from Culebra Island, Puerto Rico, **A)** male, **B)** female. *Sphaerodactylus macrolepis macrolepis* from Culebra Island, Puerto Rico **C)** adult male (with re-generated tail), **D)** adult female. *Sphaerodactylus grandisquamis guarionex* from Arecibo, Puerto Rico. **E)** male, **F)** female. Photos by ARF and RGR.

We extracted DNA from all samples using the Wizard SV^©^ Genomic DNA Purification System and associated *quick protocol* steps, and we then used the Polymerase Chain Reaction (PCR) to amplify fragments of four mitochondrial (CO1, 16S, NADH dehydrogenase subunit 2 [ND2], and cytochrome B [CYTB]) and two nuclear genes (recombinant-activating gene 1 [rag1] and oocyte maturation factor [c-mos]) encompassing a total of six genetic loci (Table S2). We visualized PCR products by gel electrophoresis and purified individual PCR products using exonuclease I/shrimp alkaline phosphatase (ExoSAP^®^). We then sequenced products in both directions on an automated sequencer (ABI 3730XL) at either the Genomic Sciences Laboratory at North Carolina State University, Raleigh, NC (nucDNA loci) or the Massachusetts General Hospital DNA Core Facility, Cambridge, MA (mtDNA loci). We assembled contigs and manually verified ambiguous base calls using Geneious^®^ 10.1.2 (Biomatters, Auckland, New Zealand). We obtained additional sequences for the above loci for an outgroup species (*S. elegans*, Cuba) and in-group (additional PRB/SXB taxa) from GenBank (data from: Hass, 1996; Gamble et al., 2008; Heinicke et al., 2011). Finally, we obtained additional CO1 sequence data for both *S. macrolepis* and *S. beattyi* from St. Croix (unpublished work, Nicole F. Angeli). Our overall dataset reflects an attempt to maximize taxon coverage and genetic locus coverage.

For each nuclear locus, we resolved heterozygous intron sequences using PHASE v. 2.1 (Stephens et al., 2001; Stephens and Donnelly, 2003) implemented in DnaSP v5.10.1 (Librado and Rozas, 2009) using default parameters for 100 iterations with a burnin of 100, and a cut-off of PP > 0.7 for base calling. We aligned our new and GenBank-mined sequences using the ClustalW 2.1 (Larkin et al., 2007) algorithm implemented in Geneious using reference sequences and default parameters. We created the following alignments for subsequent analyses: 1) a CO1 mtDNA alignment for *S. macrolepis* plus outgroups; 2) a concatenated mtDNA alignment of 16S, ND2, and CYTB for all our samples; 3) an mtDNA alignment of the ND2 locus for molecular-clock divergence-time analysis, and 4) individual alignments of both nuclear loci for gene tree/species tree analyses (Table 1).

**Table 1.** Alignments and analyses used in this study. Note that the number of sequences varies owing to the use of sequence data for some loci from GenBank or other sources. Figures where resulting phylogenetic trees are shown are listed, as are the filenames for the accessioned alignments and trees (available from https://github.com/caribbeanboas).

| Alignment | Locus (Loci) | # sequences | Total bp | Model | Analyses | Figure | Accessions |
| --- | --- | --- | --- | --- | --- | --- | --- |
| 1 | CO1 | 33 | 658 | TrN+G | ML | Fig. 4 | pr_sphaero_co1_align.nex<br>pr_sphaero_co1_tree.nex |
| 2<br>(concatenated) | 16S, ND2, CYTB | 108 | 2741 | TIM1+I+G | ML, Bayesian | Figs. S1,S2 | pr_sphaero_mtdna_align.nex<br>pr_sphaero_mtdna_MLtree.nex<br>pr_sphaero_mtdna_BStree.nex |
| 3 | ND2 | 56 | 1148 | TIM1+I+G | Bayesian | Fig. 3 | pr_sphaero_nd2_align.nex<br>pr_sphaero_nd2_beast.nex |
| 4<br>(individual) | RAG1, cmos | 44, 43 | 332, 374 | F81, JC | Bayesian<br>gene tree-species tree | Fig. S3 | pr_sphaero_rag1_align.nex<br>pr_sphaero_rag1_tree.nex<br>pr_sphaero_cmos_align.nex<br>pr_sphaero_cmos_tree.nex<br>pr_sphaero_starbeast_tree.nex |

### mtDNA Phylogenetic Analyses

We selected the best-fit model of molecular evolution for each locus (Table S2) using Bayesian information criteria (BIC) in jModelTest2 (Guindon and Gascuel, 2003; Darriba et al., 2012). We conducted Maximum Likelihood (ML) phylogenetic inference on mtDNA alignments 1 and 2 using the software RAxML (Stamatakis, 2006) via the RAxML plugin-in for Geneious. We used the GTRGAMMA model and the rapid bootstrapping algorithm with 1000 bootstrap (BS) replicates followed by the thorough ML search option with 100 independent searches. We considered BS values above 70% to indicate relatively well-supported clades (Felsenstein, 2004).

To estimate divergence times across the Puerto Rican *Sphaerodactylus* mitochondrial gene tree, we inferred a time-calibrated ND2 coalescent tree in the program BEAST v1.10 (Suchard et al., 2018) using a relaxed molecular clock model and a rate of molecular evolution of 0.65% divergence per lineage, per million years. This rate has been used previously for the ND2 locus in other lizards (Macey et al., 1998), and we note that we are primarily interested in the relative rather than absolute divergence times and thus our subsequent analyses are largely insensitive to the specific molecular clock rate used. We additionally conducted Bayesian phylogenetic analysis using BEAST on our unpartitioned concatenated mtDNA alignment implementing the Beagle library v3.0.1 (Ayres et al., 2011) to speed up computations. For both alignments (alignments 2 and 3; Table 1), we ran a MCMC for 100 million generations using a Yule speciation prior, and an uncorrelated lognormal relaxed (UCLN) molecular clock model. We repeated the analyses three times with different random number seeds, sampling every 10,000 generations and discarding the first 15% of generations as burn-in. We assured adequate mixing of the chains by calculating the effective sample size values for each model parameter, with values >200 taken to indicate adequate sampling of the posterior distribution. We assessed convergence of the independent runs by a comparison of likelihood scores and model parameter estimates in Tracer v1.5 (Rambaut et al., 2018). We combined the results from the three analyses using Logcombiner (Suchard et al., 2018) and generated a maximum clade credibility (MCC) tree with the software TreeAnotator (Suchard et al., 2018). We then estimated mtDNA genetic distances (Tamura-Nei distances) between focal clades using Mega X (Kumar et al., 2018).

We additionally conducted a gene-tree species-tree analysis for our multi-locus (ND2, 16S, CYTB, RAG1, CMOS) dataset using *BEAST (Heled and Drummond, 2010), implemented in Beast v1.8. This method estimates species tree topology under the multispecies coalescent model, which assumes that incongruence among gene trees is due to incomplete lineage sorting and not gene flow between taxa. We assigned a priori “species” designations for each recognized taxon of *Sphaerodactylus* (excluding subspecies) and treated major clades from the previous analyses as separate operational taxonomic units (OTUs). We used alignments 2 and 4 from above, reduced to include fewer individuals per OTU to improve computational time. We partitioned sequence data by locus (concatenated mtDNA and individual nucDNA loci) and assigned a locus-specific model of nucleotide substitution chosen using BIC in JModelTest2 (Table S2). We unlinked nucleotide substitution models, clock models, and gene trees in all analyses. We employed an UCLN clock model of rate variation for the mtDNA locus and a strict clock for nuclear loci (Ho et al., 2005; Peterson and Masel, 2009), and we used a Yule process speciation prior for the edge lengths. We ran the MCMC as above for 100 million generations with three independent replicates. We did not include a non-Puerto Rican outgroup as we were primarily interested in the ingroup taxa *S. macrolepis* and *S. grandisquamis* and wanted a complete data matrix. We visualized resulting trees in DensiTree v.2.2.5 (Bouckaert, 2010) and FigTree v1.4.3 (Rambaut, 2014). All alignments and trees from our phylogenetic analyses are accessioned into the online repository Dryad (doi: forthcoming).

## RESULTS

### Phylogenetic Analyses

Our concatenated mtDNA alignment included a total of 108 taxa and a maximum of 2,741 base pairs (bp) per individual, consisting of a maximum of 479 bp of 16S, 1148 bp of ND2, and 1113 bp of CYTB with 1,887 segregating sites. From this alignment we estimated both Maximum Likelihood (Fig. S1) and Bayesian (Fig. S2) phylogenies, each of which included all species of *Sphaerodactylus* described from the PRB except *S. parthenopion*. We further estimated a Bayesian molecular-clock calibrated phylogeny of the ND2 locus (Fig. 3), which included 56 taxa and a maximum of 1140 bp. All three phylogenies were topologically consistent with one another, and clearly indicated species-level divergence among all recognized Puerto Rican species. Previous proposed relationships among these species, including the lineage containing *S. klauberi, S. nicholsi, S. townsendi, S. gaigae*, and *S. monensis* (e.g., Pyron et al. 2013) were supported (Fig. 3). This clade diverged from the other major PRB clade comprised of *S. roosevelti* and *S. macrolepis* 10.2 My (95% Highest Posterior Density interval 11.87–7.98 My) (Fig. 3). Within this clade we find strong support (PP = 1; Figs. 3, S1, S2) for a divergent lineage of geckos from the PRB (with representative samples from Culebra Island, Puerto Rico and Vieques Island). This lineage is 6.74 My divergent (95% HPD 8.46–4.87 My) from its closer relative, *S. macrolepis*, and is further characterized by distinct morphological characters including male coloration (see *Morphology* below). We recognize this lineage as *S. grandisquamis* (Stejneger, 1904).

**Figure 3.**
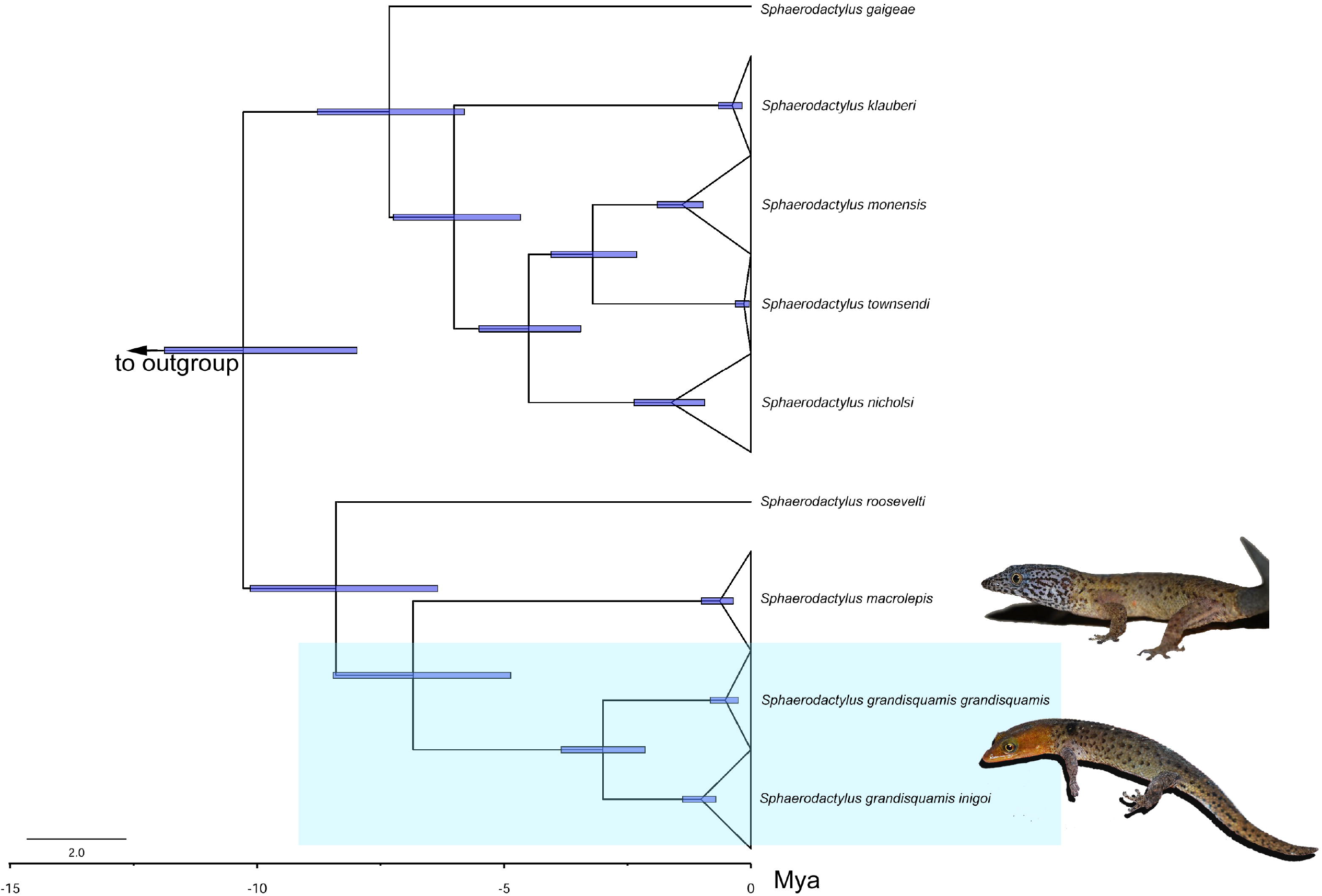
Maximum-clade credibility calibrated phylogeny for representative Puerto Rican *Sphaerodactylus*. This is a BEAST tree resulting from the analysis of the ND2 mitochondrial locus. All nodes shown are supported with posterior probability of 1. Blue bars indicate the 95% highest posterior density intervals for the coalescent times for each lineage, based on a molecular clock calibration. Images of adult male *S. grandisquamis inigoi* and *S. macrolepis* from Culebra are shown to the right.

We generated a Maximum Likelihood phylogenetic reconstruction for our CO1 alignment, which included sequence data from the topotypic *S. m. macrolepis* from St. Croix. This produced a well-resolved phylogeny that supports the *S. macrolepis* from St. Croix and Virgin Islands as conspecifics, while *S. grandisquamis* is a distinct species (BS = 99) (Fig. 4). Our gene tree/species tree approach incorporating 3,447 bp from 6 loci resolved an identical topology to the other trees, with support for distinction between *S. macrolepis* and *S. grandisquamis* (Fig. 5).

**Figure 4.**
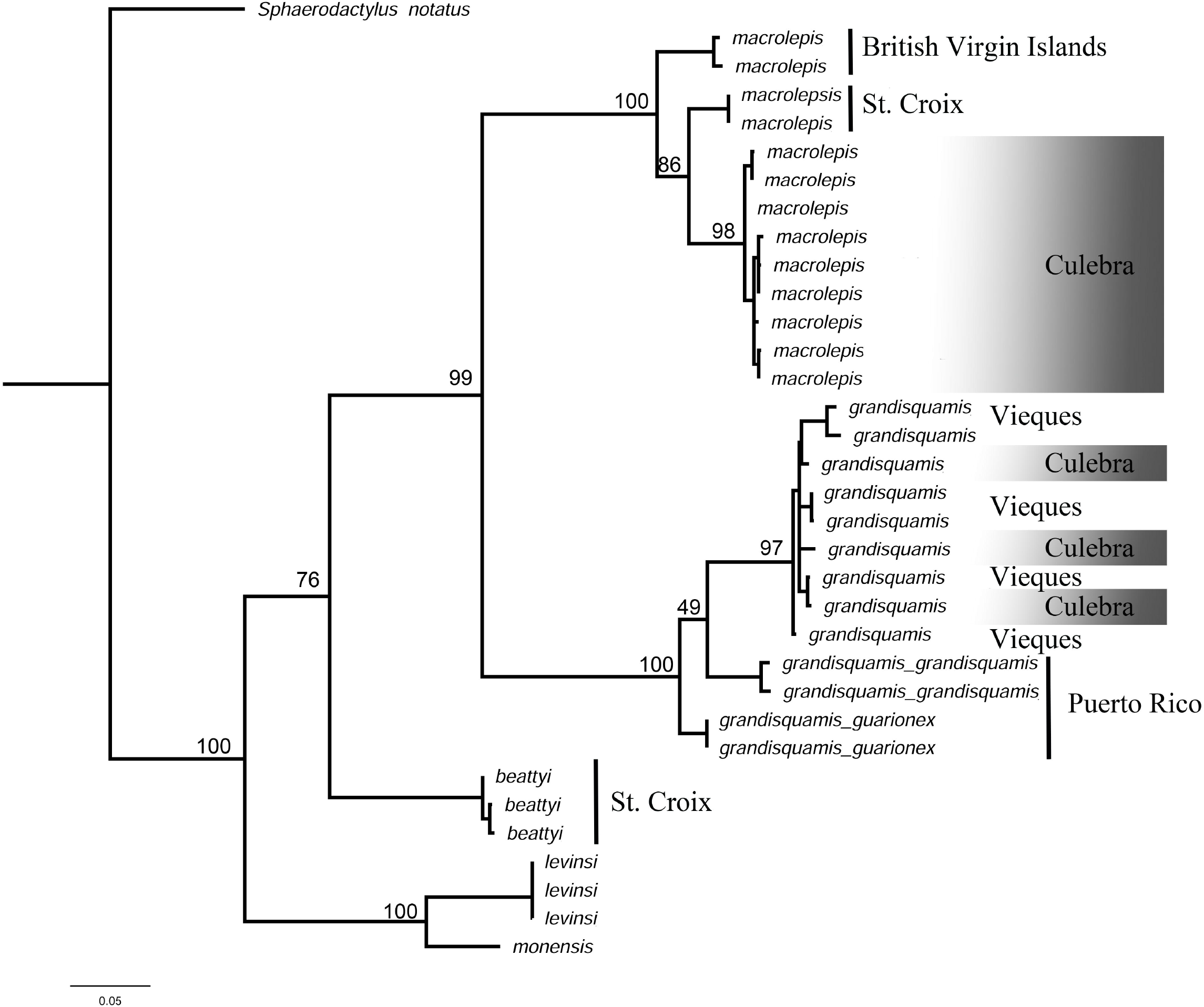
Maximum likelihood tree resulting from analysis of mitochondrial CO1 alignment. Note that topotypic St. Croix *Sphaerodactylus macrolepis* share a clade with other Culebra and Virgin Island *S. macrolepis*, while geckos from Puerto Rico, Vieques and Culebra represent the new species *S. grandisquamis*. Bootstrap support is shown for major nodes.

**Figure 5.**
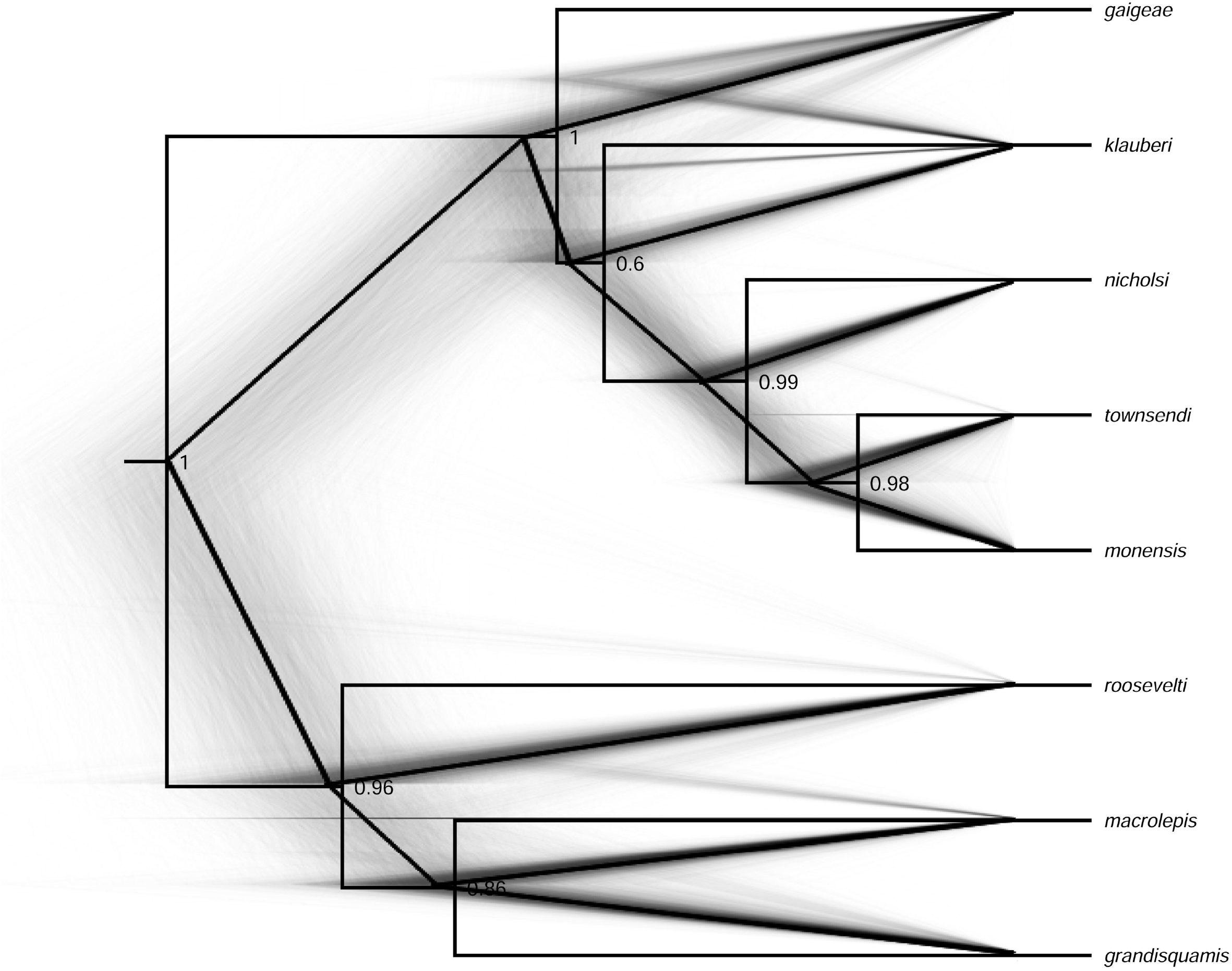
DensiTree phylogram from post-burn-in trees resulting from multi-locus gene tree/species tree analyses in *BEAST, with the maximum-clade credibility tree overlaid. These trees are generated from 3,447 bp from six loci, and numbers at nodes represent posterior probabilities (PP).

### Morphology

Our morphological assessment of Culebra geckos further supports recognition of syntopic *S. macrolepis* and *S. grandisquamis* on Culebra Island. These species, both of which are sexually dimorphic, are readily diagnosable based on coloration of both males and females (Fig. 2). They are further distinguished by a combination of meristic traits which we detail in the species re-description below.

We found that the *S. grandisquamis* from Culebra (Fig. 2A,B) are phylogenetically allied to *S. g. inigoi* from Vieques (Fig. 4; see also: Ríos-Franceschi et al. 2016). While they are not distinct populations based on the molecular data, the males from these islands show a number of unique morphological characteristics (sensu Rivero, 2008). Like Vieques lizards, Culebra *S. grandisquamis* have a tan to grayish-brown dorsal ground color, an orange to yellow throat in males and a gray throat in females, both patterned with reticulation. Culebra lizards also have a black scapular patch in males and females with small white ocelli, un-patterned heads in males and heavily striped grayish heads in females with distinct occipital and nuchal spots. Unlike the Vieques population, male Culebra animals have ample dark dorsal patterning (dark scales and/or blotches), orange heads, and orange to yellowish tails.

## DISCUSSION

Haas (1991) hypothesized that the center of the Greater Antillean *Sphaerodactylus* diversification is the island of Hispaniola, owing to the group’s relatively high species richness in that locale. Díaz-Lameiro et al. (2013) briefly discussed this, as the hypothesis is also supported by an amber-encased specimen of an extinct *Sphaerodactylus* species from Hispaniola (Daza and Bauer, 2012). Large-scale phylogenies to-date suggest that *Sphaerodactylus* on Puerto Rico are not derived from a single in-situ radiation, but instead arise from multiple ancestors originating from within the Hispaniolan radiation (Pyron et al., 2013). Our data do not directly address these higher-level phylogenetic and historical biogeographic relationships; rather, we focused on the species found on the PRB, SXB, and the islands in the Mona Passage to investigate relationships among these taxa.

There are at least 10 species of *Sphaerodactylus* distributed across the Puerto Rican Bank and islands in the Mona Passage (Mayer, 2012). Thomas and Schwartz (1966) hypothesized that ancestral states of morphological characters (“primitive characters”) among Puerto Rican *Sphaerodactylus* were best represented in the extant *S. grandisquamis*, suggesting that it might constitute the most ancestrally diverged lineage of the group. Díaz-Lameiro et al. (2013) did not find support for this hypothesis. Instead they hypothesized that *Sphaerodactylus* in Puerto Rico consisted of two major lineages: the first containing the *S. roosevelti* lineage and the *S. macrolepis* complex (*S. macrolepis* and *S. grandisquamis*), the other made up of the ecologically distinct *S. klauberi* and *S. nicholsi*. Our results also recover two clades, one containing *S. grandisquamis, S. macrolepis*, and *S. roosevelti*; the second consisting of *S. gaigeae, S. klauberi, S. monensis, S. townsendi*, and *S. nicholsi* (Fig. 3). The hypothesized relationship of rock-dwelling *S. parthenopion* with *S. nicholsi* by Thomas and Schwartz (1966) could be further tested to clarify if *S. parthenopion* represents an eastern extent to the *S. nicholsi* clade. Diaz-Lameiro et al. (2013) also hypothesized that *S. nicholsi* is potentially invalid at the specific level owing to their observation of interdigitation of *S. nicholsi* and *S. townsendi* haplotypes, although they suggest that this observation could be attributed to introgression (Murphy et al., 1984) combined with limited sampling. Pinto et al. (2019) suggest that misidentification of some samples might have been responsible for producing the pattern and recommend that specimens purported to be *S. nicholsi* and *S. townsendi* from southern Puerto Rico be re-evaluated.

### St. Croix Dwarf Geckos

Two species of dwarf geckos occur on the SXB: *S. beattyi* and *S. macrolepis*. The former is endemic to St. Croix, while the latter species also occurs on St. Croix and the PRB (Culebra Island, U.S. Virgin Islands, British Virgin Islands; Henderson and Powell, 2009; Table 2). It is unusual for a species of dwarf gecko to occur on more than one island bank in the Caribbean, although a few other cases have been reported. For example, the Cuban species *S. notatus* and *S. nigropunctatus* occur on Cuba and the Great Bahama Bank, with the former species also having colonized the Florida Bank (in the Florida Keys). Our data support *S. beattyi* as a member of the Puerto Rican *Sphaerodactylus* radiation (Fig. 4). Because the inclusion of that species is based upon a limited analysis of a single mtDNA locus (CO1) we cannot make strong inferences about its phylogenetic or biogeographic affinities relative to other species.

**Table 2.**
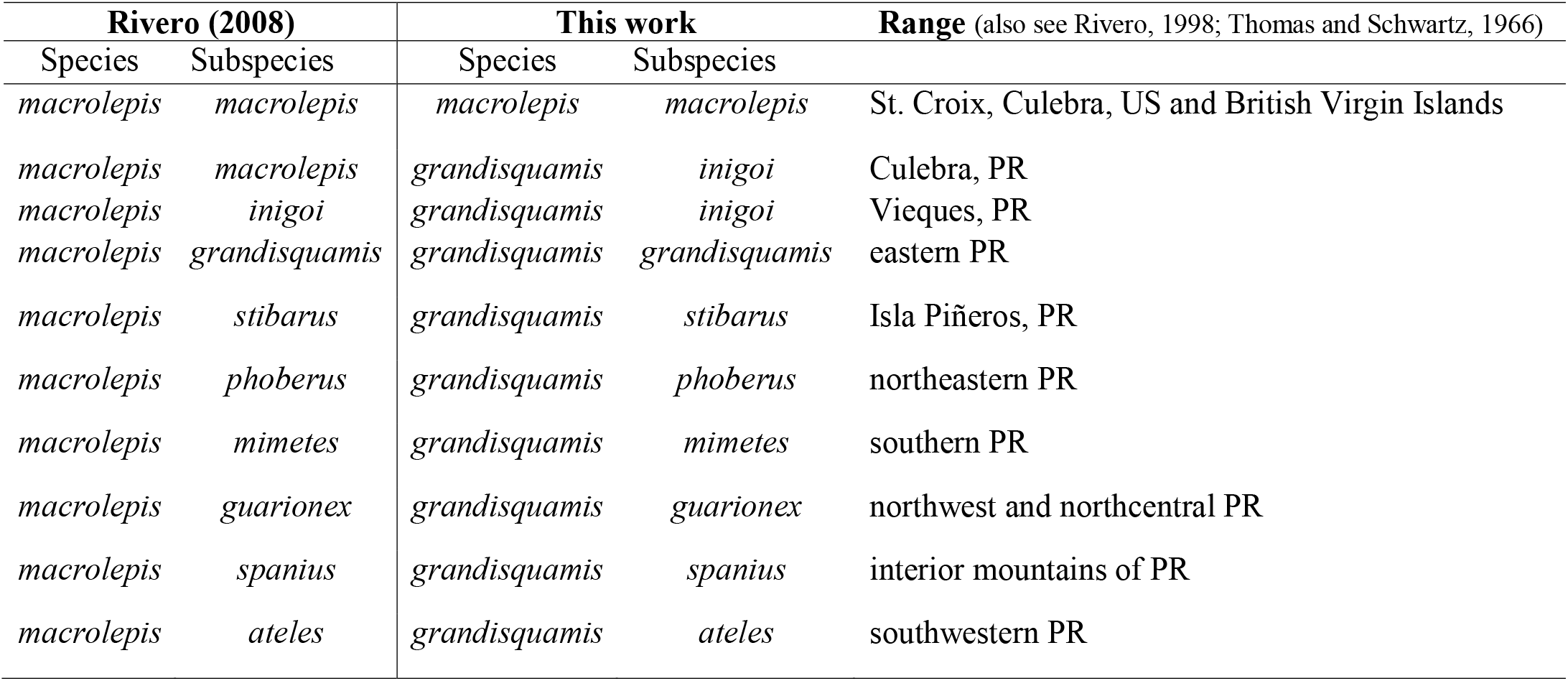
Overview of *S. macrolepis* and *S. grandisquamis* subspecies and systematic changes proposed in the present work for this group. After Schwarz and Henderson (1991).

### Sphaerodactylus grandisquamis

Our molecular data provide limited information on genetic divergence among the ten subspecies of *S. grandisquamis*, largely owing to limited sampling within each subspecies and across the geographic boundaries of these subspecies (as well as marker type). For example, we find that most subspecies represented in our dataset show little evolutionary divergence from each other (Figs. 4, S1, S2). The most apparent pattern of intraspecific divergence that we found is that *Sphaerodactylus grandisquamis* subspecies from the main island of Puerto Rico are highly divergent (PP =1, BS = 100, 95% HPD = 3.84–2.15 My) from the Culebra and Vieques subspecies (*S. g. inigoi*). Within the island of Puerto Rico, *Sphaerodactylus g. grandisquamis* are not well-differentiated from *S. g. guarionex* (PP = 0.92, BS < 50; Figs. S1, S2). *Sphaerodactylus g. spanius* appears to be sister to *S. g. guarionex* in our ML tree (BS = 98; Fig. S1) although not in our BS tree (PP<0.90; Fig. S2). While several consistent morphological traits appear to diagnose subspecies (Thomas and Schwartz, 1966; Rivero 1998; this work), future researchers interested in this question might consider taking a fine-scale morphological and population genetic approach to characterizing diversity and divergence of this widespread taxon.

### Culebra Island Geckos

Culebra Island is small but topographically complex and is situated in a region of biogeographic interest both as an emergent portion of the PRB but also as a link between the mesic habitats of mainland Puerto Rico to the west and the more xeric habitats of the Virgin Islands to the east. Prior to the present study, a single species of dwarf gecko was recognized from Culebra: *S. macrolepis*. This widespread species was thought to consist of nine subspecies on the Puerto Rico Bank (Thomas and Schwartz, 1966; Rivero, 1998; Schwartz and Henderson, 1991; Table 2). A tenth subspecies, *S. m. parvus*, was described from the Leeward Antilles (King, 1962) but has since been elevated to *S. parvus* (Powell and Henderson, 2001). *Sphaerodactylus macrolepis* on Culebra was originally described as the species *S. danforthi* (Grant, 1931a,b), although it was synonymized with *S. m. macrolepis*, the same subspecies as the topotypical population on the St. Croix Bank (Thomas and Schwartz, 1966). One of the distinguishing characteristics of this species is the pronounced sexual dichromatism, with males having a distinctly grayish head with darker stippling or reticulation (Thomas and Schwartz, 1966; Fig. 2C,D).

During a focused herpetofaunal survey of the Monte Resaca region on the island of Culebra, a Spanish Virgin Island found approximately 10 km to the east of the Puerto Rican mainland (Fig. 1), Ríos-Franceschi et al. (2016) collected specimens of typical *S. macrolepis* coloration (Fig. 2C,D). They also observed what appeared to be two additional morphological types of dwarf geckos that seemed to be distinct in shape and coloration from *S. m. macrolepis* and *S. m. inigoi*, suggesting that more than one species or more than two subspecies occurred on Culebra and its satellite island of Luis Peña. One of these geckos we tentatively identified as *S. townsendi* (Fig. S4), although we were not able to obtain a tissue sample to verify this identification. It has been suggested that *Spherodactylus townsendi* occurs on Culebra (Rivero, 1998, 2006; Mayer, 2012). However, if it ever was present on that island, it was subsequently no longer thought to occur there (Henderson and Powell, 2009; Joglar et al., 2017; but see Fig. S4).

Multi-locus Bayesian and ML analyses of up to 4,104 bp of DNA sequence data (maximum of 1,887 mtDNA and 49 nucDNA segregating sites) of the remaining two groups found on Culebra revealed that *S. m. macrolepis* (Fig. 2C,D) occurs on the island of Culebra, along with the newly-elevated species *S. grandisquamis* (Fig. 2A,B). *Sphaerodactylus grandisquamis* is also found on islands to the south and east (Vieques, Puerto Rico, and satellite islands [Fig. 1]).

### Species concept interpretation

We follow the General Lineage Species Concept (de Queiroz, 1998) in conjunction with the Evolutionary Species Concept (Wiley, 1978) with regard to our definition of species. We use molecular phylogenetics and distinguishing sets of morphological characteristics to support our hypothesis and identification. We find strong supportive evidence for the recognition of *S. grandisquamis* (Stejneger, 1904)— a new species of dwarf gecko from the Puerto Rico Bank. Our definition is based on consistent strong support in Maximum Likelihood and Bayesian mtDNA phylogenetic reconstructions, an estimated divergence time of 6.74 My (95% HPD 8.46–4.87 My) from *S. macrolepis*, 10.8 % sequence divergence from syntopic *S. macrolepis*, and distinguishing morphological traits detailed below. Importantly, *S. grandisquamis* and *S. macrolepis* are syntopic and yet genetically distinct.

## SYSTEMATIC ACCOUNT

*Sphaerodactylus grandisquamis* (Stejneger, 1904)

(Figure 2 A,B,E,F)

ZooBank registration: urn:lsid:zoobank.org:act:52C0978E-1C17-4FA8-B83B-18D3AE30E58E

### *Sphaerodactylus macrolepis* Gunther *1859*

#### Holotype

USNM 27007, adult male collected by L. Stejneger on 4th March 1900 at Luquillo, Puerto Rico.

#### Distribution

This type locality of this species is Luquillo, Puerto Rico. It is distributed across the main island of Puerto Rico, some satellite islands, Culebra Island and satellites, and Vieques Island and satellites. It does not occur east of the Spanish (=Passage) Islands.

#### Etymology

The specific epithet is an elevation of the previously-recognized subspecies *S. macrolepis grandisquamis* (Thomas and Schwartz, 1966). The epithet is Latin meaning “possessing large scales.”

#### Diagnosis

*Sphaerodactylus grandisquamis* is a medium-sized (34–35mm SVL), sexually dichromatic species that can be distinguished from other Puerto Rico Bank (and close relatives) congeneric geckos via the following characters: from *Sphaerodactylus gaigeae* Grant 1932 by non-cycloid ventrals, sexual dichromatism (versus the non-sexually dichromatic *S. gaigeae*), absence of chevron-like marks near the nape or scapular area (versus 2 anterior chevron-like patterning near head and neck in *S. gaigeae*); from *S. klauberi* Grant 1931 by smaller body size (SVL >36 mm in *S. klauberi*), scales weakly to moderately keeled (versus strongly keeled in *S. klauberi*), fewer midbody scales (31–54 versus 42–67 in *S. klauberi*), sexually dichromatic (versus absence of sexual dichromatism in *S. klauberi*); from *S. levinsi* Heatwole 1968, sexually dichromatic (absence of sexual dichromatism in *S. levinsi*), found on main island of Puerto Rico (*S. levinsi* is restricted to Isla Desecheo); from *S. monensis* Meerwarth 1901 chest scales non-keeled (versus presence of keeled chest scales in *S. monensis*), throat rarely immaculate (versus immaculate pinkish grey in *S. monensis*), found on main island of Puerto Rico (*S. levinsi* is restricted to Isla Desecheo); from *S. nicholsi* Grant 1931a, non-keeled ventral chest scales (versus presence of occasional chest scale keeling in *S. nicholsi*), sexually dichromatic (versus absence of sexual dichromatism in *S. nicholsi*), absence of a postocular stripe extending into tail (present in *S. nicholsi*); from *S. roosevelti* Grant 1931a slightly smaller size (*S. roosevelti* around 39 mm), heavily sexually dichromatic (versus weakly sexually dichromatic in *S. roosevelti*); from *S. townsendi* Grant 1931a larger body size (SVL 28 mm in *S. townsendi*), sexually dichromatic (versus absence of sexual dichromatism in *S. townsendi*), lacking distinct black sacral stripes (usually present in *S. townsendi*; Fig. S4); from *S. micropithecus* Schwartz 1977 sexually dichromatic (versus absence of sexual dichromatism in *S. micropithecus*), found on main island of Puerto Rico (*S. micropithecus* is restricted to Isla Monito). The above characters and descriptions follow from Grant (1931, 1932), Thomas and Schwartz (1966), Schwartz and Henderson (1991), and Rivero (1998, 2006).

*Sphaerodactylus grandisquamis* substantially differs genetically from other PRB *Sphaerodactylus* with a mtDNA pairwise Tamura-Nei sequence divergence of 9.9%–13.2% (Table S3).

#### Similar Species

Syntopic *Sphaerodactylus macrolepis* (Günther, 1859; Fig. 2C,D) on Culebra Island have the following characteristics: gular region speckled with khaki and black granular scales; throat coloration same as gular; ventral surface of forelimbs pale-khaki colored; dorsal surface of forelimbs tan to brownish; ventral surface of tail slightly darker than ventral surface of abdomen, lightly speckled; dorsal surface of hind limbs tan, slightly more speckled than forelimbs; ventral surface of hind limbs; “salt-and-pepper” head patterning; 18 axilla-groin scales; 34–35 scales around the midbody; male dorsum dark tan or khaki-colored with scattered darker scales; absent of a scapular patch; an SVL of 22–23 mm; three supralabial scales and 3 infralabial scales; nine 4^th^ toe lamellae; seven 4^th^ finger lamellae; and gular scales smooth, many granular.

#### Natural History

This species is a leaf-litter denizen, foraging in the top layers of leaf litter under forest shade as a presumably dietary generalist feeding on small arthropods such as termites (Steinberg et al., 2007; ARF pers. ob.).

#### Taxonomic Notes

This species contains eight recognized subspecies (Table 2).

## Acknowledgements

We are grateful to the Puerto Rico Departamento de Recursos Naturales y Ambientales (DRNA) and the Virgin Islands Department of Planning and Natural Resources (DPNR) for permits and assistance. Novel samples were collected under DRNA permit 2013-IC-007 (to RGR) and DPNR permit DFW16052U (to NFA). We acknowledge support from the University of Massachusetts Boston, Harvard University, the Harvard Museum of Comparative Zoology, the University of North Carolina Asheville, the University of Puerto Rico Mayagüez, and the Global Genome Initiative under Grant No. GGI-Rolling-2015-PRBVI. Portions of this work have been approved by the University of Massachusetts Boston Institutional Animal Care and Use Committee (IACUC) Protocol #2012001 and Smithsonian Institution Animal Study 2017-06. We are grateful for tissue specimen loans from the Museum of Vertebrate Zoology, University of California Berkeley; and the Yale Peabody Museum, Yale University. We especially thank Dan Mulcahy for curatorial information regarding some samples, and George Zug for reviewing an earlier version of this manuscript.

## Data Accessibility

Associated data archived at Figshare (DOI: 10.6084/m9.figshare.11372079) and GitHub (https://github.com/caribbeanboas)

## Appendix 1.

### Supplemental Tables

**Table S1.** Specimens and localities for tissue samples used in this study.

Download from Figshare (DOI: 10.6084/m9.figshare.11372079) and GitHub (https://github.com/caribbeanboas)

**Table S2.**
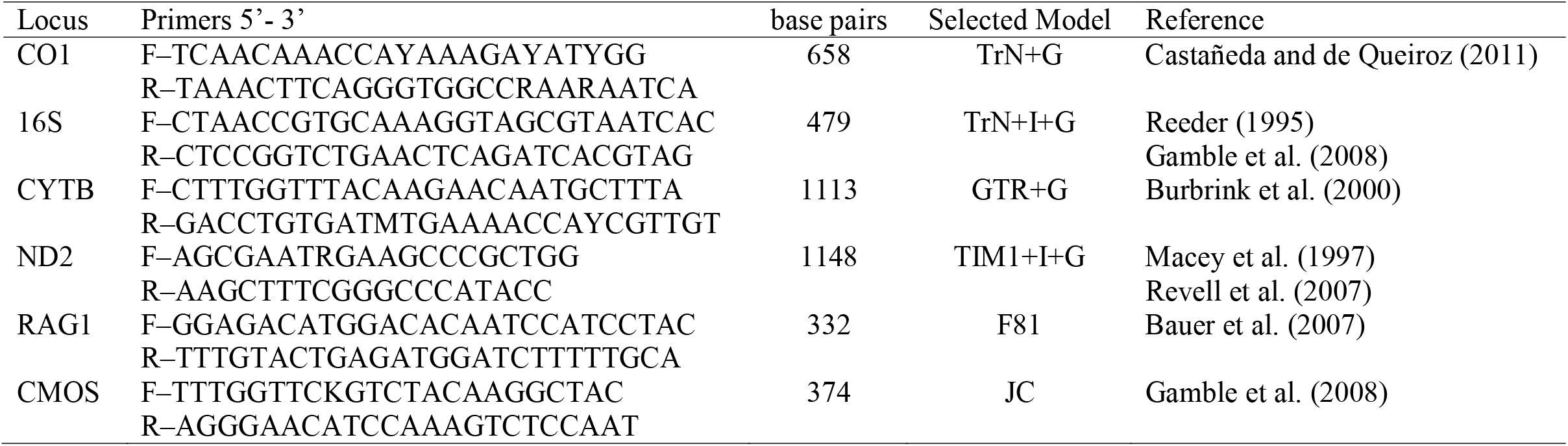
Primers and loci used in this study.

**Table S3.** Mean genetic distances between *Sphaerodactylus* species on the Puerto Rico Bank. Distances are net Tamura-Nei distances measured from a 2,741 bp alignment of three mtDNA loci (16S, CYTB, and ND2).

|  | *S. grandisquamis* | *S. macrolepis* | *S. nicholsi* | *S. townsendi* |
| --- | --- | --- | --- | --- |
| *S. grandisquamis* | – |  |  |  |
| *S. macrolepis* | 0.108 | – |  |  |
| *S. nicholsi* | 0.110 | 0.090 | – |  |
| *S. townsendi* | 0.099 | 0.091 | 0.019 | – |
| *S. klauberi* | 0.132 | 0.129 | 0.075 | 0.073 |

## Appendix 2.

### Supplemental Figures

High resolution figures can be downloaded from Figshare (DOI: 10.6084/m9.figshare.11372079) and GitHub (https://github.com/caribbeanboas)

**Fig. S1.** Maximum likelihood tree resulting from analysis of three concatenated mitochondrial loci (16S, CYTB, and ND2).

<u>Download here</u>

pr_sphaero_mtdna_MLtree.nex

pr_sphaero_mtdna_MLtree.pdf

**Fig. S2.** Bayesian tree resulting from analysis of three concatenated mitochondrial loci (16S, CYTB, and ND2).

<u>Download here</u>

pr_sphaero_mtdna_BStree.nex

pr_sphaero_mtdna_BStree.pdf

**Fig. S3.** Maximum likelihood trees resulting from the analysis of two concatenated nuclear genes (cmos and RAG1).

<u>Download here</u>

pr_sphaero_rag1_tree.nex

pr_sphaero_cmos_tree.nex

**Figure S4.**
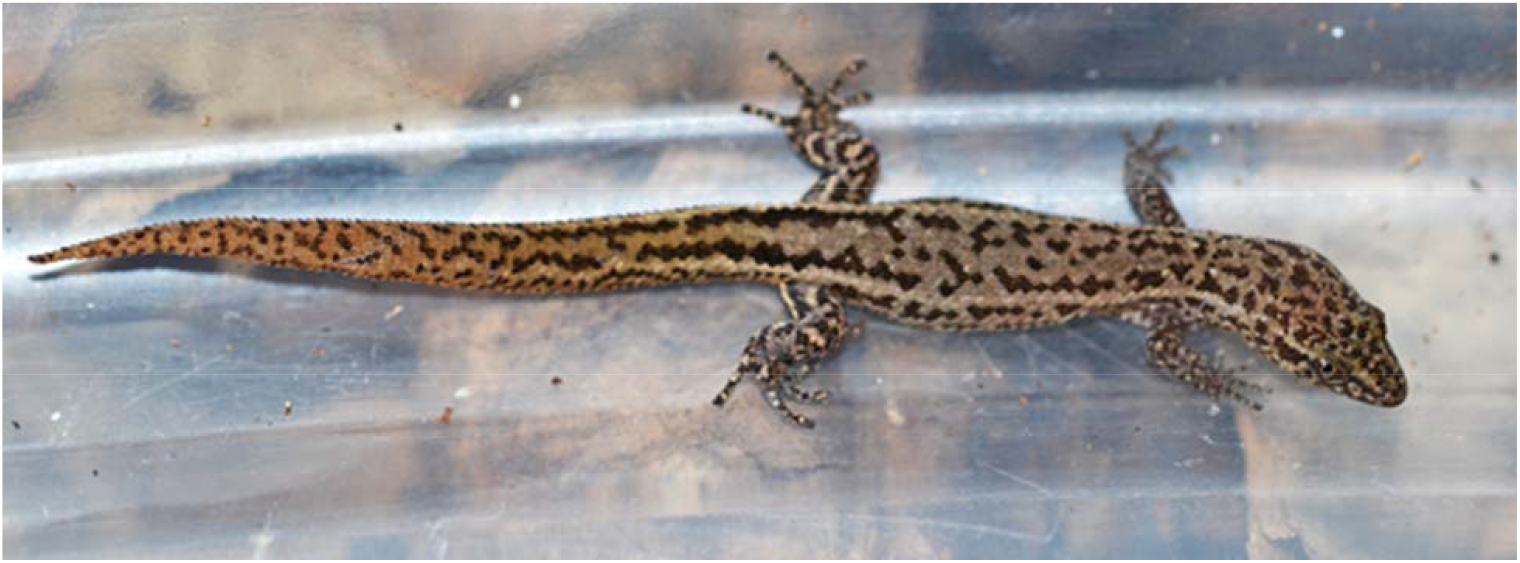
*Sphaerodactylus cf. townsendi* from Culebra Island. Photo by ARPR.

